# Muscle Stem Cell Niche Dysregulation in Volumetric Muscle Loss Injury

**DOI:** 10.1101/346395

**Authors:** Shannon E. Anderson, Woojin M. Han, Vunya Srinivasa, Mahir Mohiuddin, Marissa A. Ruehle, Austin Moon, Eunjung Shin, Cheryl L. San Emeterio, Molly E. Ogle, Edward A. Botchwey, Nick J. Willett, Young C. Jang

## Abstract

Skeletal muscle has a remarkable regenerative capacity; however, after volumetric muscle loss (VML) due to traumatic injury or surgery this regenerative response is significantly diminished, causing chronic functional deficits. The critical defect size at which the muscle will not functionally recover has not yet been established and subsequently, the relative contribution of crucial muscle components, including muscle stem cells and the muscle stem cell niche, are unknown. In this study, we created VML injuries of 2, 3, or 4 mm diameter, full-thickness defects in the mouse quadriceps. The 2, 3, and 4 mm injuries resulted in a defect of 5, 15, or 30% of the quadriceps mass, respectively. At 14 and 28 days after injury, histological analyses revealed injury size-dependent differences in myofiber morphology and fibrosis; the number of small myofibers increased with increasing injury size. The results showed that the 3 mm injury was at a threshold point, as myofibers were unable to bridge the defect, there was persistent fibrosis and inflammation, and significantly increased number of myofibers with centrally located nuclei. We then further investigated the 3 mm VML for nerve and vascular regeneration. These injured muscles were accompanied by a drastic increase in denervated neuromuscular junctions (NMJ), while assessment of angiogenesis via micro-CT analysis revealed a significant increase in vascular volume primarily from small diameter vessels after VML injury. Collectively, these data indicate that the spatial and temporal control of the fibrotic and neuromotor response are critical to regeneration and could be potential therapeutic targets, as they are the most dysregulated components of the muscle stem cell niche after VML.

## Introduction

Skeletal muscle has a remarkable endogenous regenerative capacity when the basal lamina remains intact (Mackey & Kjaer, 2017; Webster et al., 2016). In the most commonly investigated experimental skeletal muscle injury models, such as freeze injury, barium chloride, notexin, and cardiotoxin, muscle is capable of full regeneration with virtually no fibrosis (Hardy et al., 2016; Pollot & Corona, 2016). However, within the field of musculoskeletal injuries, volumetric muscle loss (VML) remains a persistent challenge. VML injuries are defined as a traumatic or surgical loss of tissue which will not endogenously regenerate, resulting in chronic loss of function (Grogan & Hsu, 2011). VML is most commonly seen clinically as the result of extremity trauma injuries, which account for more than 50% of all trauma injuries among both military and civilian populations (Banerjee et al., 2013; Dougherty et al., 2009; Owens et al., 2008). As VML injuries result in a chronic loss of muscle function, these injuries result in permanent disability (Corona et al., 2015). The current gold standard of care for VML injuries is autologous free or rotational muscle flap transfer; while these are successful procedures for limb salvage, these techniques remain largely unsuccessful for regenerating of functional muscle tissue and additionally cause donor-site complications and morbidity (Garg et al., 2015; Grogan & Hsu, 2011).

One major factor which sets VML injuries from other common skeletal muscle injury models is the catastrophic loss of the basal lamina as well as all other supporting cells (Grogan & Hsu, 2011; Pollot & Corona, 2016). These biophysical and biochemical components make up the specific microenvironment for muscle stem cells (MuSCs), also called satellite cells. Situated between the basal lamina and the myofiber sarcolemma, MuSCs are influenced by a unique microenvironment, termed the MuSC niche. This niche is comprised not only of the adjacent myofiber and extracellular matrix (ECM), but also the closely associated vasculature, interstitial stromal cells, such as fibro/adipogenic progenitors (FAPs), and neural networks such as motor neurons, neuromuscular junction, and terminal Schwann cells. These cellular and acellular niche components work cooperatively at different stages of successful myofiber regeneration, and are needed for the formation of functional skeletal muscle and maintenance of muscle homeostasis (Joe et al., 2010; Yin et al., 2013). MuSCs are indispensable for skeletal muscle regeneration, as the ablation of this cell population will result in a total loss of functional muscle regeneration (Lepper, Partridge, & Fan, 2011). Due to the striking differences in functional outcomes between muscle injury models where this niche remains and those where the niche components are entirely removed we hypothesize non-healing VML injuries reach a critical size due to a significant dysregulation in the regenerative response of the endogenous supporting ECM, vasculature, and neuromuscular innervation. A characterization of VML in this way will fill a current gap in knowledge as there is currently limited understanding of at what point the muscle stem cell niche is critically dysregulated, as well as a lack of knowledge about which stem cell niche components are critical for functional recovery.

In researching VML injuries, it is necessary to establish animal models which recapitulate commonly seen clinical phenotypes. Clinically, VML injuries are categorized as total compartment loss, where the nerve supplying the injured compartment is lost, or partial compartment loss, where the supplying nerve remains. The injury location which most often leads to high rates of requested late amputation are those to the lower extremities. Additionally, above-knee VML is typically more severe and has more limited treatment options (Grogan & Hsu, 2011), however, in pre-clinical animal models, below knee VML models, such as in the tibialis anterior (TA), have been more commonly studied (Pollot & Corona, 2016). In this study, to best replicate the critical VML that is commonly observed clinically, we used a pre-clinical mouse model of VML in the quadriceps with varying sizes. The small rodent models are useful for elucidating the underlying cellular pathology after VML due to the availability of transgenic models and histological reagents. By choosing a full thickness defect affecting all four constituent muscles in the quadriceps, we can more accurately recapitulate the severity and location of clinically manifested VML injuries (Li et al., 2014; Willett et al., 2016). Additionally, the threshold for a “critical size” for VML injury models has not yet been established; whereas this is commonplace in other large volume defect models in other tissue types, namely bone (Roddy, DeBaun, Daoud-Gray, Yang, & Gardner, 2018). The resulting inconsistencies in size and location of defects in VML research makes it difficult to directly compare across studies (Grasman et al., 2015). The characterization of a pre-clinical model which recapitulates aspects of clinical phenotypes will be helpful in the design of therapeutic strategies.

In this manuscript we set out to define the “critical size” in a VML defect and characterize the key components that are dysregulated at this threshold. To achieve this, we evaluated multiple sizes of VML defects to determine the defect size where there is ineffective myofiber bridging of the defect due to the presence of an unresolved fibrotic and inflammatory response. We defined these parameters by investigating the response of key muscle stem cell niche components including the extracellular matrix and fibrotic deposition, macrophage presence, neuromuscular re-innervation, and re-vascularization. This comprehensive characterization of muscle stem cell niche dysregulation in non-healing skeletal muscle injuries will ultimately serve to inform the next generation of therapeutic targets in the treatment of VML injuries.

## Materials and Methods

### Animals

C57BL/6J mice were purchased from Jackson Laboratories. Wild-type mice were used for all experiments with the exception of experiments assessing neuromotor regeneration, for which B6.Cg-Tg(Thy1-YFP)16Jrs/J (Jackson Laboratories #003709) mice were used. Thy1-YFP constitutively express yellow fluorescent protein (YFP) via the Thy1 gene promoter, which is expressed in all motor neurons. Adult female mice between 3-9 months in age were used for all experiments. All animals were housed and treated according to the protocols approved by the Georgia Institute of Technology Institutional Animal Care and Use Committee.

### Volumetric Muscle Loss Injury

Mice were anesthetized from inhalation of 1-3% isoflurane. Sustained-release buprenorphine (0.8 mg/ml) was administered for pain management. The incision site on the hindlimb was shaved and hair removal lotion was used to remove remaining hair. The area was disinfected using ethanol and chlorohexidine wipes. A unilateral incision was made to expose the quadriceps muscle. Biopsy punch tools of 2, 3, or 4 mm (VWR, 21909-132, −136, −140) in diameter were used to make consistent full-thickness defects to the mid-belly region through all four constituent muscles of the quadriceps (the rectus femoris, vastus lateralis, vastus intermedius, and vastus medialis). After injury, the skin was closed using wound clips. For wet weight analyses, 36 mice were used, 12 for each time point: 7, 14, or 28 days. For each timepoint, there were three different injury sizes, 2, 3, or 4 mm muscle biopsy, and there was n=4 animals per defect size. These same mice were used for histological analyses, at the 14 and 28 day time points. For neuromuscular junction imaging, 8 Thy1-YFP mice were used total, all of which received 3 mm defects, with 4 mice used for each time point, either 14 or 28 days. For microCT angiography, 5 mice received 3 mm defects and used for 28 day time points. At the appropriate time point, animals were euthanized by CO_2_ inhalation.

### Tissue Wet Weight Analysis

After creating the injury, the skin was closed, and animals were left to heal for 7, 14, or 28 days without intervention. Immediately after euthanasia, the animals were weighed and the hindlimb muscles were dissected. The wet weight of the muscles was obtained and was normalized to the total body weight of each animal.

### Tissue histology and immunostaining

Quadriceps muscles were dissected from the hindlimbs of mice and stored on ice cold PBS until wet weight was obtained. Muscles were then snap frozen in liquid nitrogen cooled isopentane and stored at −80°C. Cyrosections were taken on a CryoStar NX70 Cryostat at 10 µm thickness. Hematoxylin and Eosin staining (VWR, 95057-844, −848) was performed. Gomori’s Trichrome staining kit (Polysciences, 24205-1) was used according to the kit instructions. Prior to antibody staining, tissue sections were blocked and permeabilized using blocking buffer (5% BSA, 0.5% goat serum, 0.5% Triton-X in 1X PBS) for 30 minutes at room temperature and then blocked with Goat F(ab) Anti-Mouse IgG (Abcam, ab6668) diluted to 2 µg/ml in blocking buffer for 1 hour if anti-mouse secondary antibodies were necessary. Samples were washed in between each step with 0.1% Triton in PBS. Primary antibodies were diluted in blocking buffer at a ratio of 1:200 for dystrophin (Abcam, ab15277), vWF (Abcam, ab6994), and CD68 (Abcam, ab125212). Primary antibody for embryonic myosin heavy chain (eMHC) (DSHB, F1.652) was diluted 1:10 in blocking buffer. All primary antibodies were incubated for 1 hour at room temperature. Secondary antibodies were conjugated to Alexa Fluor 488 (Thermo Fisher, Ms: A-11029, Rb: A-11034), 555 (Thermo Fisher, Ms: A-21424, Rb: A-21429), or 647 (Thermo Fisher, Ms: A-21236, Rb: A-21245). All secondary antibodies were diluted 1:250 in blocking buffer and allowed to incubate for 30 minutes at room temperature. Alexa Fluor 647-conjugated phalloidin (Thermo Fisher, A22287) was diluted 1:250 in blocking buffer and incubated with secondary antibodies for 30 minutes at room temperature. Slides were mounted with Fluoroshield Mounting Medium with DAPI (Abcam, ab104139) and stored at 4°C.

### Tissue preparation for neuromuscular visualization

Hindlimbs were harvested above the hip joint and fixed in 4% paraformaldehyde for 1 hour at room temperature. Hindlimbs were stored at 4°C in PBS until processing. Both quadriceps muscles were dissected from the mouse hind limbs and placed in blocking buffer (5% BSA, 0.5% goat serum, 0.5% Triton-X in 1X PBS) for 1 hour. The dissected muscle was then stained with α-Bungarotoxin conjugated with Alexa Fluor 647 (Thermo Fisher, B35450, 1:250 in PBS) and anti-GFP conjugated with Alexa Fluor 488 (Novus Biologicals, NB600310X, 1:200 in PBS) for 30 minutes at room temperature. The tissue within the defect site was then cut into smaller segments. Muscle segments were whole mounted on glass slides with Fluoroshield Mounting Medium with DAPI (Abcam, ab104139) and stored at 4°C.

### Tissue preparation for microCT angiography

Immediately after euthanasia with CO_2_ gas, mouse vasculature was perfused with saline until the liquid ran clear, approximately 30 mL of total solution. The perfusion line was switched to 10% Neutral Buffered formalin for an additional 30 mL, or until tissue was completely fixed. Finally, tissue was perfused with Microfil contrast agent (Flow Tech, MV-122, 1:2 mix with diluent) until vessels were visibly perfused. Tissue was stored at 4°C until dissection.

### Second Harmonic Generation Imaging of collagen

Images were taken on a Zeiss 710 Laser Scanning Confocal Microscope using 20X objective and stitched together with Zen Black software (Zeiss) to create a complete image of the entire cross-section. Second harmonic generation was achieved using a Pulsed IR laser at 810 nm with the confocal pinhole entirely opened. Signal from CD68 antibody, conjugated with Alexa Fluor 555 secondary antibody was imaged simultaneously.

### Confocal imaging and quantification of fiber cross-sectional area and eMHC^+^ fibers

Immunofluorescence images were taken on either a Zeiss 700 or Zeiss 710 Laser Scanning Confocal microscopes at 20x and stitched together with Zen Black software (Zeiss) to create a complete image of the cross-section. The fluorescence channel corresponding to the dystrophin images was analyzed using ImageJ. Images were converted to 8-bit, thresholded, and muscle fiber cross sectional area was measured through the Analyze Particle function for particles of 0.25-1.0 circularity and 150-6000 µm^2^ in area. The ImageJ output would overlay a mask on each fiber it measured, giving it a number corresponding to the measurement taken. Area was output directly in ImageJ measurements and these values were used to create a histogram in GraphPad Prism 7. Data was grouped by injury size or contralateral control, with 3 replicates taken from an n=4 animals for each group. The histogram displays the relative frequency of counts in each bin as a cumulative histogram, with bin size equal to 100 µm^2^. Quantification of eNMC^+^ myofibers was done on images containing all channels (eNMC, dystrophin, DAPI). The brightest fibers were counted using the ImageJ multipoint tool. Each replicate slide was counted twice and all counts for each injury size were analyzed.

### Imaging and quantification of neuromuscular junctions

Z-stack images were taken on a Zeiss 710 Laser Scanning Confocal using 40X objective. Z-stacks of 5 random fields of view were taken from whole mounted sections from within the injured area or from a comparable region from the contralateral control. NMJs were quantified by eye by a blinded scorer (Tse et al., 2014). In this way, the number of NMJs were quantified by placement in one of three categories. The categories were considered (1) normal, pretzel-like morphology, (2) fragmented, or abnormal morphology, or (3) newly forming AChR clusters (Jang et al., 2010).

### MicroCT angiography imaging and analysis

Quadriceps were dissected from fixed, Microfil perfused mice. The muscle was wrapped in radio-transparent foam for microcomputed tomography (microCT) imaging in a vivaCT 40 (Scanco Medical) at 21 µm voxel size. All scans were performed with an applied electrical potential of 55 kVp and current of 145 µA. Analysis was done using native Scanco software suite for all muscles on the middle section of the muscle, in the area where the VML injury was made. The analyzed volume of interest (VOI) was a cylinder with a diameter of 6.429 mm and a height of 250 slices, or a total length of 5.25 mm. A voxel density threshold of 105 was applied for segmentation of vasculature, and a Gaussian filter was used to suppress noise (σ = 1.2, support = 2). The native analysis scripts within Scanco software suite was used to generate the measurements for perfused vascular volume in the analyzed area, as well as vessel diameter distribution (incremental 21µm bins).

### Statistical Analysis

All statistical analyses were done in GraphPad Prism 7. Data shown as mean +/− standard error of the mean (S.E.M.) for all figures. One-way ANOVA was used to analyze defect wet weight (Fig. 1B) and eMHC^+^ fiber quantification (Fig. S1A), two-way ANOVA was used to analyze the wet weight of injured quadriceps over time (Fig. 1C) and centrally located nuclei (Fig. 4J), with both using Tukey’s test as a *post-hoc.* For each time point, the cumulative histograms for each injury size and control group were all compared to one another using multiple Kolmogorov-Smirnov tests. Paired, two-tailed *t*-tests were done to analyze microCT data (Fig. 6C,D, Fig. S3).

**Figure 1.**
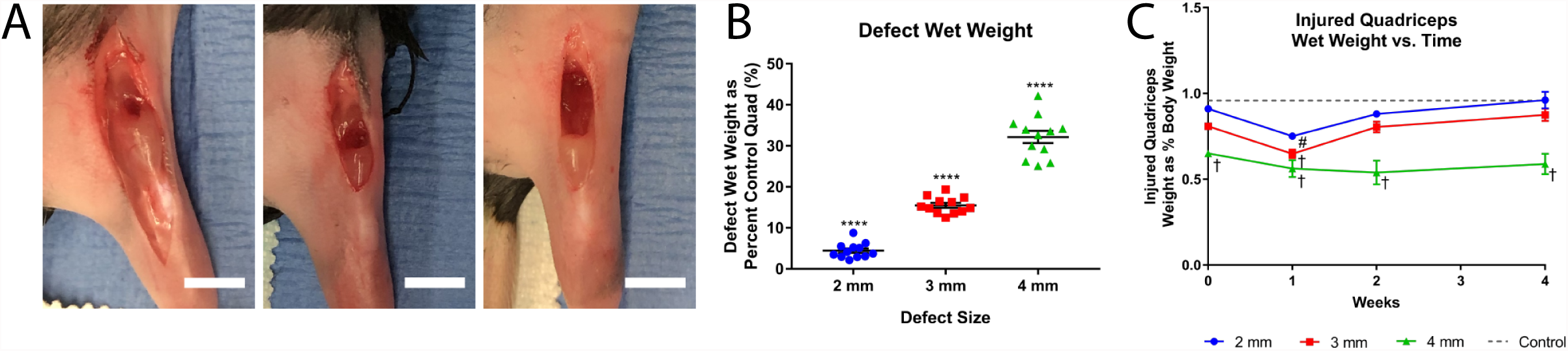
Various sized VML injuries in mouse quadriceps. (A) Mouse quadriceps after removal of biopsied muscle tissue. From left to right, 2 mm, 3 mm, and 4 mm biopsy punches. Scale bars represent 5 mm. (B) The wet weights of biopsied quadriceps muscle normalized to the contralateral control as plotted for each biopsy punch size. Mean percentage of contralateral control is 4.44%, 15.5%, and 32.2% for 2, 3, and 4 mm injuries, respectively. (n=12 per injury size, error bars indicate mean +/− standard error of the mean (SEM), (**** = *p*<0.0001 after one-way ANOVA and Tukey’s test). (C) Wet weight of the injured quadriceps 7, 14, and 28 days post injury normalized to total body weight. Day 0 values are calculated as an average of the defect wet weight subtracted from its respective Day 7, 14, and 28 contralateral control quadriceps wet weight and normalized to the body weight of the animal, mean is plotted +/− SEM. The dotted line is the average value of contralateral control quadriceps normalized to body weight. (n= 4 for each group, # = *p*<0.01, † = *p*<0.0001 as compared to the control after two-way ANOVA and Tukey’s test *post-hoc*).

## Results

### Muscle mass following full-thickness VML model in mouse quadriceps

To determine a critical size for VML injuries in the mouse hindlimb, we created defects of multiple sizes in the mouse quadriceps muscle. Biopsy punches of either 2, 3, or 4 mm were used to make full-thickness defects through the mouse quadriceps (Fig.1 A). These injuries resulted in the removal of 4.44 ± 1.85%, 15.49 ± 2.04%, and 32.16 ± 5.14% of the total quadriceps wet weight, respectively, as compared to the contralateral control (Fig. 1B). The mass loss of each of these injuries were significantly different from one another (n= 12, *p*<0.0001). Compared to the average percentage of total body weight of the contralateral control muscles, all injury sizes caused a significant decrease (n=4, 2 mm, p<0.01, 3 and 4 mm, *p*<0.0001) in quadriceps wet weight for the experimental groups after 7 days (Fig. 1 C). After 14 or 28 days, however, only the 4 mm injury group remained significantly different (n=4, *p*<0.0001) from the contralateral control.

### Qualitative histomorphology after VML injury of multiple sizes

Histomorphology was used to determine the type of tissue infiltration into each VML defect site. Cross-sections were stained with hematoxylin and eosin (H&E) to examine the general histomorphology of each injury size at 14 (Fig. 2 A) and 28 (Fig. 2 B) days post-injury, using 4 replicates for each injury size and time point. Contralateral control samples from both 14 and 28 days post-injury showed normal, healthy skeletal muscle tissue. These samples showed densely packed myofibers with peripherally located myonuclei and few other cell types. Qualitative observation of the injured samples showed vastly different morphology and composition. 2 mm injuries at both time points showed myofibers with centrally located nuclei surrounding areas of small, white pockets which resemble areas of fatty infiltration. Additionally, there were areas surrounding these pockets where mononuclear cells were embedded in a dense matrix. As the size of the injury increased, so did the alteration of the muscle appearance. In 3 mm injuries at 14 days, there were more regenerating muscle fibers, indicated by the increased number of small, centrally nucleated myofibers compared to 2 mm injuries. There was also an increase in the area where non-muscle cell types were present in between individual myofibers. At 28 days, these differences were even more apparent in 3 mm injuries. The clearest damage was in 4 mm injuries after both time points. There were large areas where no myofibers could be seen; in these regions there were dense clusters of mononuclear cells with no clear tissue structure in addition to fatty infiltration. Around the area of damage, there were newly regenerating fibers with centrally located nuclei and small cross-sectional area.

**Figure 2.**
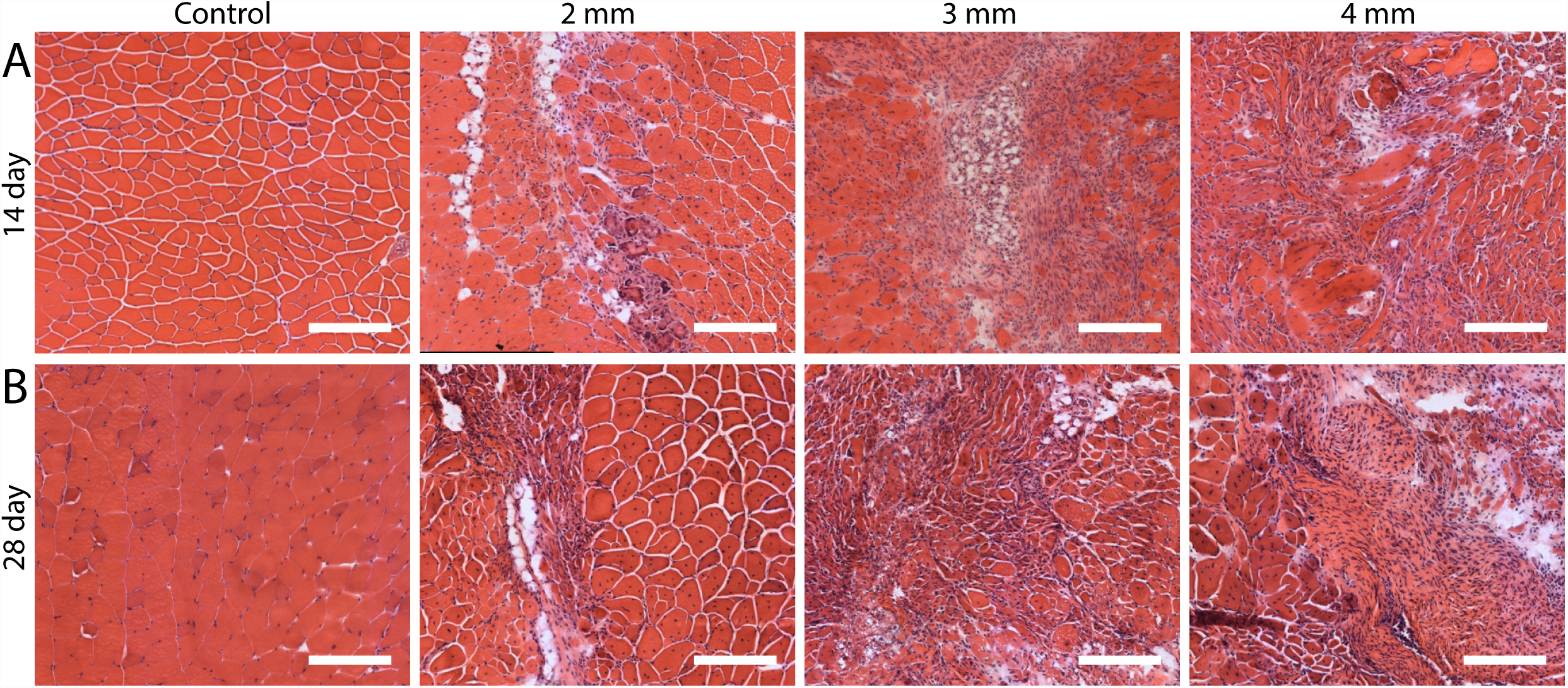
H&E staining of each injury size at various time points. Images of quadriceps cross-sections, stained with H&E. From left to right are representative images of a contralateral control, 2mm injury, 3mm injury, and 4mm injury, from 14 day (A) and 28 day (B) time points. Scale bars represent 200 µm.

### Assessment of fibrosis in multiple VML injury sizes

To determine the extent of fibrosis and matrix deposition in each VML injury size consecutive slides were stained with Gomori’s Trichrome for colorimetric imaging of collagen and imaged with Second Harmonic Generation (SHG) microscopy to visualize collagen fluorescence (Fig. 3). The slides used for SHG imaging of collagen were also stained for the pan-macrophage marker CD68. The same tissue samples used here were also used for H&E imaging, with 4 replicates at each injury size and time point. The contralateral control sections at both 14 and 28 day time points (Fig. 3A,B) showed little to no collagen signal, outside of the expected amount present in the myofiber ECM, in both Gomori’s and SHG imaging. Additionally, there were few CD68+ macrophages present in the contralateral control tissue. In contrast, the VML injured quadriceps, showed extensive collagen fluorescence and qualitatively more infiltration of CD68^+^ macrophages. 2 mm injuries (Fig. 3C,D) showed localized collagen deposition and macrophage infiltration at 14 days, while the surrounding myofibers remained largely unaffected. After 28 days, the localized fibrosis remained, however there were very few macrophages remaining. By comparison, 3mm injuries after both 14 and 28 days (Fig. 3E,F) showed qualitatively more fibrosis, seen from both SHG collagen imaging as well as Trichrome staining. Additionally, there was qualitatively more infiltration and persistence of CD68+ macrophages out to 28 days in the 3 mm VML samples as compared to 2 mm samples or the contralateral controls. In the 4 mm VML injury animals, features of fibrosis were most prominent (Fig. 3G,H). A large region of collagen fluorescence was seen at both 14 and 28 days post-injury, with many CD68^+^ macrophages present within this tissue at both time points. With increased VML injury size, the fibrotic response increased as did the persistence of unresolved macrophages. While it was clear from histomorphology that VML injuries of 3 and 4 mm diameter had substantial deposition of non-muscle tissue in the defect region, there were still many small myofibers around the defect with centrally located nuclei, which are both indicative of myofiber regeneration.

**Figure 3.**
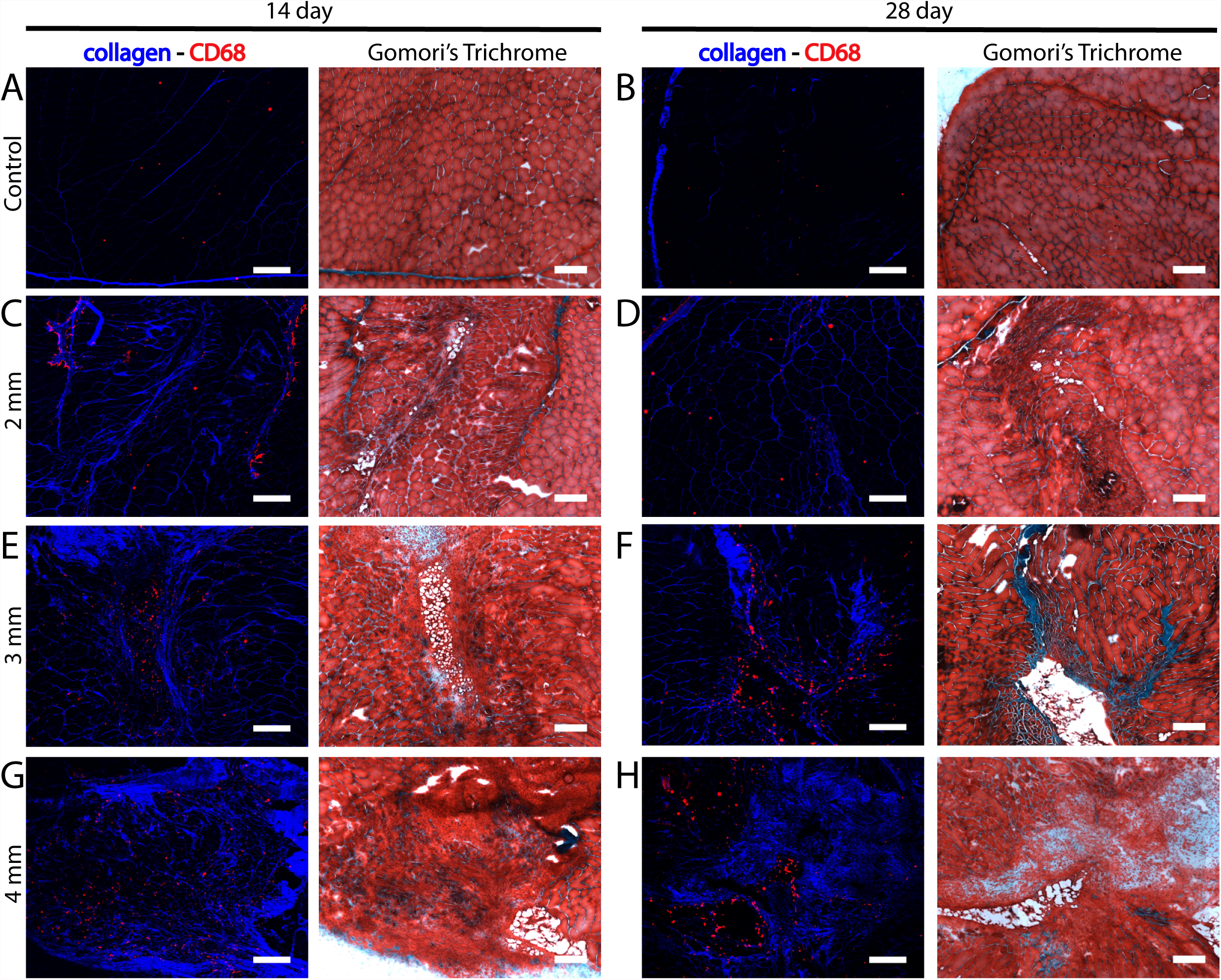
Assessment of fibrotic response at each injury size. Quadriceps cross-sections 14 (A, C, E, G) or 28 (B, D, F, H) day time points post-injury. For each set of images (A-H) the left-hand is an image of a cross-section stained with CD68 antibody (shown in white) and imaged on a 2 photon scanning confocal microscope (at 810 nm) for second harmonic generation imaging of collagen (shown in blue) and the right-hand image is colorimetric image of a cross-section stained with Gormori’s Trichrome. The sets of images are representative sections from contralateral control (A,B), 2mm injury (C,D), 3mm injury (E,F), or 4mm injury (G.H) samples. Scale bars represent 200 µm.

### Myofiber regeneration in multiple VML injury sizes

To evaluate myofiber regeneration, we quantified the size distribution of the myofibers present in 2, 3, and 4 mm VML injuries. Cross-sections were immunostained for the myofiber membrane protein dystrophin, the intracellular protein present in newly formed myofibers embryonic myosin heavy chain (eMHC), and the nuclear stain DAPI (Fig. 4A-G). Sections from the contralateral controls (Fig. 4A) showed mature myofibers with peripherally located myonuclei. The signal from the dystrophin channel was used to quantify myofiber cross-sectional area. Measured cross-sectional areas were analyzed using a histogram to determine trends of myofiber cross sectional area across injury size and time point. Both cumulative (Fig. 4H,I) and relative (Fig. S1A) histograms were generated. At both 14 (Fig. 4H) and 28 (Fig. 4I) days post-VML the data showed that 4 mm injuries had the largest relative number of myofibers with small cross-sectional areas, as the 4 mm line had the greatest slope, increasing quickly over a small number of bins. The relative accumulation rates in the cumulative histograms decreased with decreasing injury size; indicating that as injury size decreased the number of myofibers with smaller cross-sectional areas also decreased. Control cross-sections had the lowest relative number of small myofibers, which was expected, as small fiber cross-sectional area indicates regenerating myofibers. Descriptive statistics, including mean, median, and standard error, for the cross-sectional area data displayed in these histograms showed differences in distribution of values across injury size (Table S1). This data similarly illustrated that fiber cross-sectional decreased, on average, with increasing injury size at both 14 and 28 day timepoints. For both 14 and 28 days, the cumulative distributions of each injury size and the control were determined to be significantly different from all other groups via multiple Kolmogorov-Smirnov tests (*p*<0.0001). These results indicate that there are an increasing number of regenerating myofibers with increasing injury size, as regenerating myofibers will have smaller cross-sectional areas.

**Figure 4.**
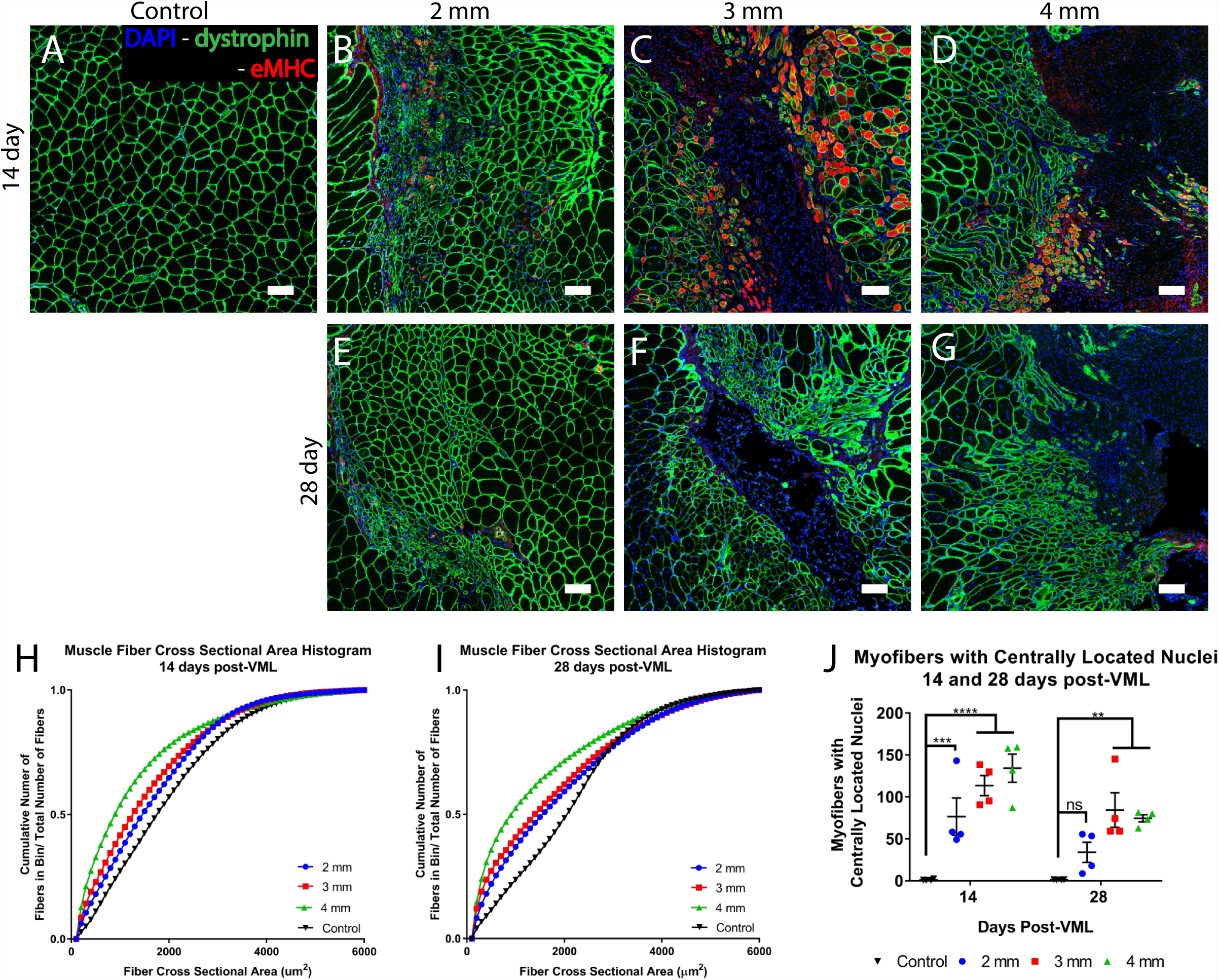
Visualization and quantification of fiber cross-sectional area in each injury size. Representative immunofluorescence images of each time point and injury size: (A) contralateral control, (B, C, D) 14 day time point of 2mm, 3mm, and 4mm injuries, respectively, and (E, F, G) 28 day time point of 2mm, 3mm, and 4mm injuries, respectively. Each of the sections were stained for dystrophin (green), DAPI (blue), and eMHC (red). Scale bars represent 100µm. Cross-sectional area was quantified using stitched images of the entire cross-section from the dystrophin channel. The measured cross-sectional areas used to create cumulative histograms 14 (H) and 28 (I) day time points. Data was separated by injury size. Each category (control, 2, 3, & 4mm) had n=4 individual animals with 3 replicate slides per animal, one replicate from the beginning, one from the middle, and one from the end of the damaged area. In the case of the control, replicate slides were taken from corresponding areas to the damaged samples. eMHC+ fibers were quantified at 14 days post-injury (J). Five representative areas from each of the same replicate slides used for cross-sectional area quantification were used to count centrally located nuclei per area. All data points shown, with bars representing mean +/− SEM. Two-ways ANOVA performed with Tukey’s test *post-hoc*, significance for *p*<0.05.

As cross-sectional area indicated that there were differences in myofiber regeneration between injury sizes at both 14 and 28 days post-VML, we assessed this further with quantification of myofibers which were embryonic myosin heavy chain (eMHC) positive and had centrally located nuclei. eMHC is protein which is transiently expressed during the early stages of myofiber development (Schiaffino, Rossi, Smerdu, Leinwand, & Reggiani, 2015), and therefore myofibers which stain positively for eMHC can be classified as newly regenerated. While new myofibers are expressing eMHC, they will also have centrally located nuclei, however the presence of centrally located nuclei will remain after eMHC is no longer expressed. These indicators can therefore give temporal information about where the myofibers are in the process of regeneration. The number of eMHC^+^ myofibers were quantified in all VML injury sizes at 14 days post-injury (Fig. S1C). Due to variability between animals, there were no significant differences in the number of eMHC^+^ fibers between any of the injury sizes. At 28 days, however, there were essentially no eMHC^+^ fibers in any defect size, indicating that there are no fibers early in the regeneration process at this time point. However, at all time points and in all injury sizes there were myofibers with centrally located myonuclei, indicating that regeneration had been recently occurring. The number of centrally located nuclei per area was also quantified (Fig. 4J). 14 days post-VML there were significant differences in the number of centrally located nuclei in all injury groups as compared to the contralateral control. By 28 days after injury, there were significant differences only between the 3 and 4 mm injury groups compared to the contralateral control leaving non-significant differences between 2 mm injuries and the contralateral control (n=4, significance for *p*<0.05). These results indicate that there are large populations of proliferating and differentiating myogenic precursor populations after VML in all injury sizes after 14 days. However, at 28 days, 2 mm injuries have regenerated to a point where there are no longer more centrally located fibers in than in the contralateral control tissue.

The results from the study comparing different VML defect sizes indicated that a 3 mm injury was at the threshold for a non-healing critical size at the 28 day time point. These injuries displayed a significant increase in sustained attempted muscle regeneration as indicated by the elevated number of myofibers with centrally located nuclei. Even with these sustained attempts, 3 mm injuries still maintained a substantial amount of unresolved fibrosis which the regenerating muscle was unable to penetrate. The persistent fibrosis was evident from the trichrome and SHG collagen imaging. Additionally, 3 mm injuries at 28 days still had persistent presence of macrophages (CD68^+^ cell population), indicating unresolved inflammation. Each of these findings contributed to our determination that 2 mm injuries (4.44 ± 1.85%) were subcritical defects, 4 mm injuries (32.16 ± 5.14%) were critical defects, and that 3 mm injuries (15.49 ± 2.04%) represent a transition point from sub-critical to critically sized. This transition point warranted further investigation into both the neuromuscular and vascular response into the injury space.

### Neuromuscular regeneration after critical VML

We investigated changes to additional muscle stem cell niche components, starting with neuromuscular regeneration, at the 3 mm threshold to a critical size injury to create a more comprehensive picture. A 3 mm diameter VML defect was made in the quadriceps of eight Thy1-YFP mice to assess neuromuscular recovery at 14 and 28 days (n=4 per time point). Neuromuscular junctions (NMJs) were quantified to determine the level of denervation or re-innervation into the injury space.

Muscle segments were stained with fluorophore-conjugated α-Bungarotoxin (BTX) to visualize acetylcholine receptors (AChRs) on the myofibers. This channel (Fig. 5 C-F) was used to quantify the number of NMJs which fit each of the 3 categories: (1) normal, pretzel-like, (2) abnormal, fragmented morphology, and (3) newly formed AChR clusters. In contralateral control samples, there was an overall higher number of NMJs (Fig. 5A), 100% of which were classified as normal, innervated NMJs (Fig. 5B) as is apparent from the representative maximum intensity projection images (Fig. 5C-F). In contrast, there were far fewer NMJs of any kind in the VML injured muscles (Fig. 5A). Of the NMJs present in VML injured muscle at 14 days post injury, 88.8% displayed fragmented morphology and therefore in group 2 and 12.5% were in group 3 as they were newly forming AChR clusters. At 28 days post-VML the NMJs present in injured muscle were 57.6% classified as group 2 and 42.4% classified as group 3. However, there were no normal, innervated NMJs at either time point into the injury site. Representative images of injured tissue at 14 (Fig. 5D) and 28 (Fig. 5F) days are qualitatively analogous, both with some fragmented (group 2) NMJs. Additionally, in these samples there was substantial amount of autofluorescence in the YFP/GFP channel (Fig. S2A-D), likely due to pervasive fibrotic tissue. VML injured tissue samples also showed some newly regenerated AChR clusters (Fig. S2E, indicated by orange arrowheads), however there was no evidence of a regenerating nerve towards these newly formed junctions, even at 28 days. Consistent with our eMHC^+^ data which showed regenerating myofibers with centrally located nuclei, there are clearly regenerating fibers following a normal regenerative program and developing AChR receptor clusters for innervation. We see no evidence, however, of final maturation and innervation of these fibers, indicating that they would remain non-functional.

**Figure 5.**
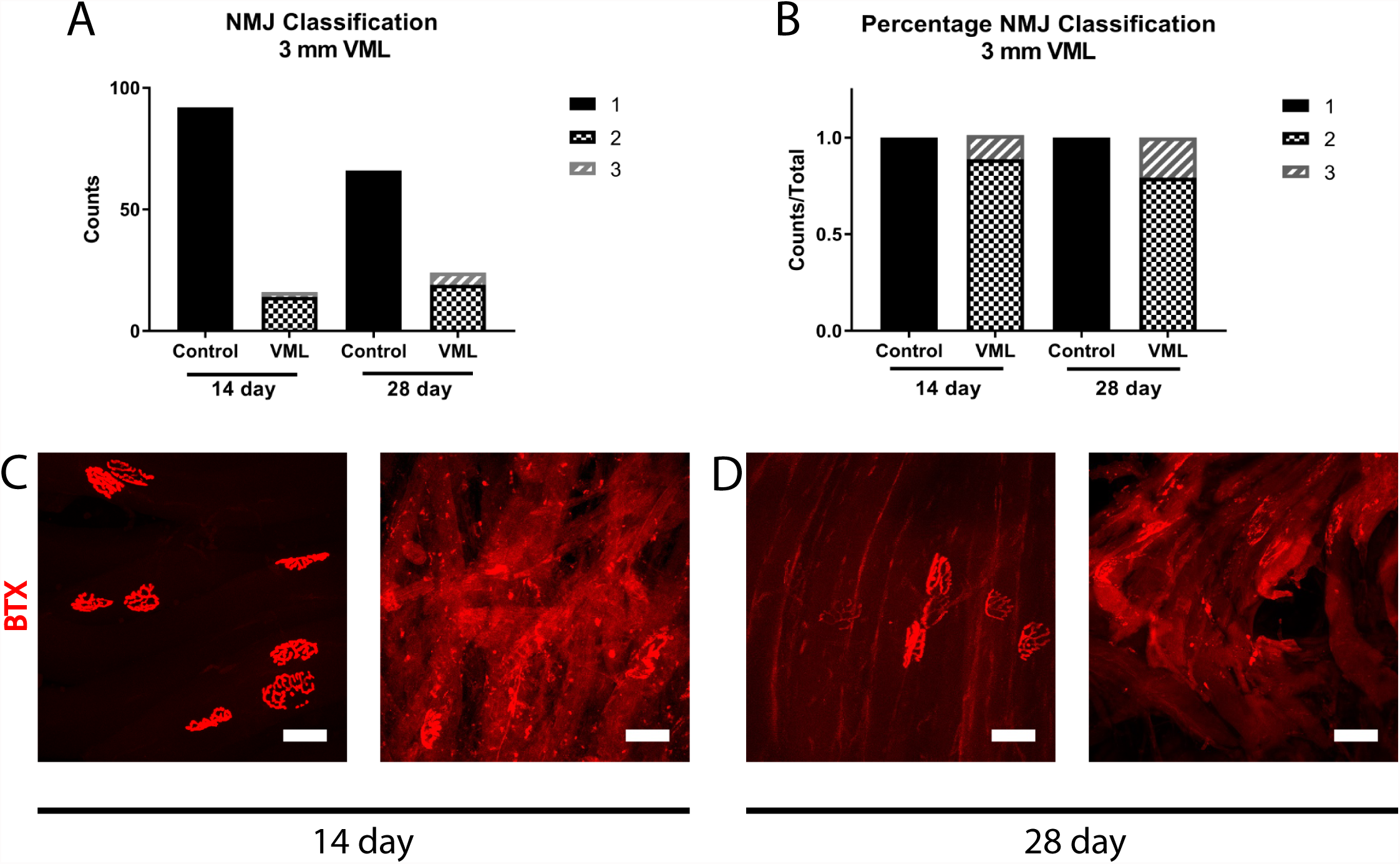
Whole mounted sections of neuromuscular junctions in VML compared to the contralateral control. NMJs were quantified by three classifications (A,B) as either group 1, 2, or 3 in Thy1-YFP mice 14 or 28 days post 3 mm VML injury. Group 1 NMJs were normal, pretzel-like morphology, group 2 NMJs were abnormal, fragmented morphology, and group 3 were newly forming AChR clusters. (A) Shows raw count data for each experimental group and time point. (B) Depicts the same data as in (A), but displayed as a fraction of total number of NMJs in each category. Representative maximum intensity projections of z-stacks taken of the control and injured experimental groups are shown for 14 (C,D) and 28 (E,F) day time points. Each image shows only the BTX channel from the image to visualize the morphology of the post-synaptic AChRs. The BTX channel was used for quantification. All scale bars are 50 µm.

### Vascularization after critical VML injury

Another key component of the muscle stem cell niche is the dense vascular network, necessary to provide nutrients to functionally active skeletal muscle myofibers. For this reason, adequate vascularization into the defect area is crucial after VML. To determine the amount of vascularization that occurs after a critically sized VML injury, 5 mice were given 3 mm VML injuries. 28 days post injury, the vasculature of each animal was perfused with Microfil, a lead contrast agent. MicroCT scans were done on each dissected muscle (Fig.6A,B), and then the middle third of the quadriceps (the area of injury) was chosen for analysis. We quantified total vascular volume within each quadriceps (Fig. 6C), allowing for pairwise comparison between each injured quadriceps and its contralateral control. This data showed significantly greater perfused vascular volume in the injured quadriceps when compared to the uninjured control. In addition, there were significant differences in the diameter of perfused vessels of the injured muscle compared to its contralateral control (Fig. 6D). The number of vessels in each diameter bin were compared pairwise with their contralateral control. These analyses (Fig. S3A-H) indicated significant increases in the number of vessels within the injured quadriceps with diameters of 21, 42, 105, 126, and 147µm. This increase in vascularization at 28 days was confirmed with immunostaining muscle cross-sections for von Willebrand factor (vWF) (Fig. 6E). Images of control and injured cross-sections show increased vascular network in the injury space. These vessels also had an increased diameter compared to those seen in the contralateral control muscle. This is consistent with the data from microCT angiography.

**Figure 6.**
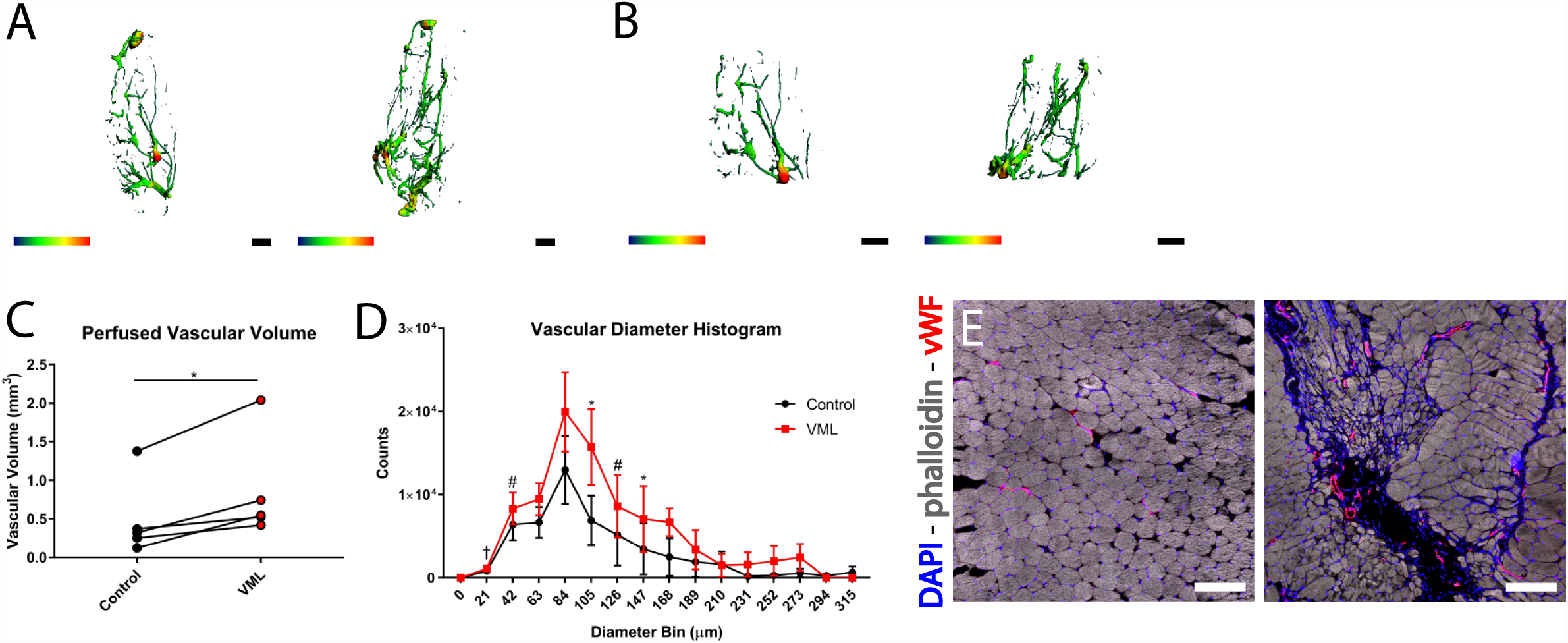
MicroCT and IHC analysis of vasculature in VML compared to control quadriceps. (A) Reconstructed 3D heat map of µCT images from microfil perfused contralateral control and 3mm injured quadriceps, left to right, from the same animal. (B) Reconstructed 3D heat map of the middle third of the same samples as in (A). The middle third of each sample is what was quantitatively analyzed in (C) and (D). (C) Total perfused vascular volume for the middle third of each sample shown in a pairwise comparison to match injured and control from the same animal. (D) Histogram counting the number of vessels in each diameter bin. The bins represent the resolution of the measurement itself (e.g. vessels between 0 and 21 µm are placed in the 21 µm bin). Counts are shown as mean +/− SEM from 5 samples, significance was determined via a paired, two-tailed t-test. Color scale is from 0.000 to 0.315 mm for vascular diameter, length scale bar is 1 mm. Paired, two-tailed t test performed for vascular volume comparison and for comparison of injured and control values in each bin, * *p*<0.05, # *p*<0.01, † *p*<0.0001. Pairwise comparisons from values in each bin shown in Supplementary Data. (E) Images taken of quadriceps cross-sections 28 days after a 3 mm injury (right) and its contralateral control (left). Staining was done for phalloidin(gray), nuclei (blue), and vWF (red). Positive vWF staining shows large diameter vessels within the defect space.

## Discussion

The goal of this study was to provide general guidelines on the critical size threshold in a full-thickness animal model of VML. For our full-thickness quadriceps defect model in mice, we determined the transition to a critically-sized defect was a biopsy punch 3 mm in diameter, as myofibers were unable to bridge the defect space in an injury of this size. These 3 mm injuries constituted a loss of approximately 15% of the muscle mass. Previously it has been cited that a 20% muscle loss is the threshold for failure of the native regenerative process (Turner & Badylak, 2012), however this number is not well supported in the literature. This non-healing threshold was determined in the context of MuSC niche dysregulation. As the MuSC is essential for muscle regeneration (Lepper et al., 2011), it is crucial that its niche components, including the ECM, vasculature, and motor neuron innervation, properly modulate the microenvironment to direct successful muscle regeneration. At the critical threshold, there is a chronically increased fibrotic and inflammatory response, shown by increased collagen deposition and CD68^+^ cell populations after 28 days post-VML. Additionally, there is no evidence of re-innervation of any newly regenerating myofibers as well as an increased number of large diameter vessels in the injury space as compared to uninjured contralateral muscle.

The most similar study to determine a critical sized muscle injury was done nearly 20 years ago, testing the gap length that muscle could regenerate into after pre-clinical rat model of muscle transection, or lesion (Terada, Takayama, Yamada, & Seki, 2001). This study isolated a bundle of transected muscle fibers, *in vivo* using a silicone tube, and separated the ends by a known distance. In their study, they show that isolated muscle fibers are capable of only bridging a 1 mm gap, filling the 2 mm gaps with fibrotic tissue, and a 3 mm gap remains entirely vacant with muscle ends being capped by fibrotic scar tissue. This injury model was inherently different from ours, but allows us to draw parallels to our full thickness defect model as they previously highlighted the phenomenon of a fibrotic cap covering the ends of all severed muscle fibers creating a clear physical barrier to regeneration. Therefore, in our multiple sized full-thickness defect analysis in an environment that more closely replicates clinically seen phenotypes, the results indicate surrounding muscle and support structures can allow for bridging of a 2 mm defect. We see, however, that the 3 mm defect created a critically sized gap where a fibrotic barrier prevented bridging and healthy skeletal muscle regeneration.

We have shown that there is a distinct phenotype as a result of a critically sized defect. The presence and persistence of collagen fibrosis and fatty infiltrate in the environment increased with injury size. This fibrotic and fatty infiltrate response is characteristic of skeletal muscle trauma, as has been shown consistently in previous VML studies (M. T. A. Li et al., 2014; Matthias et al., 2018; Sicari et al., 2012). Additionally, similar results have been seen in various ECM scaffold tissue engineered strategies, where the scaffolds themselves become populated with fibrotic tissue comparable to an empty defect (Corona & Greising, 2016; Garg et al., 2014; Greising et al., 2017). While this fibrotic tissue is generally considered one of the main barriers to successful muscle regeneration, it is also important to note that complete ablation of muscle resident fibroblasts leads to altered regenerative capacity of satellite cells (Murphy, Lawson, Mathew, Hutcheson, & Kardon, 2011), indicating that when properly regulated, fibroblasts are a vital niche component in muscle regeneration. Therefore, determining a method for modulating this fibrotic response to be pro-regenerative will be essential in VML therapeutics.

In addition to verification of this previously well-characterized phenotype following muscle trauma, our study also identified additional phenotypic markers for a critically sized VML defect. While macrophage infiltration has been studied in the short term during the typical inflammatory period after VML (San Emeterio, Olingy, Chu, & Botchwey, 2017), our results show a clear persistence of CD68^+^ macrophages even after 28 days post-injury. This phenotype sets a VML injury apart from many other acute injury types and draws similarities to chronic muscle disorders, such as muscular dystrophy (Wallace & McNally, 2009; Wehling-Henricks et al., 2010). In these such disorders, chronic muscle damage results in the deposition of fibrotic tissue and fatty infiltrate between myofibers, resulting in chronic functional deficits, similar to VML injuries. The resulting chronic inflammatory phenotype has been shown to result in a unique macrophage phenotype which promotes the sustained proliferation of FAPs (Lemos et al., 2015), which could potentially be a driving force behind the persistent and dysregulated fibrosis in VML injuries as well. This would indicate potential for therapeutic interventions targeting this chronic inflammation for creating a pro-regenerative microenvironment post-VML.

Additionally, we observed a distinct temporal pattern in myofiber regeneration pattern after VML. eMHC expression was present 14 days after injury, which is unexpected after acute muscle injury. Typically, in more commonly characterized types of skeletal muscle injury, eMHC expression is not seen after 7 days post-injury (Elabd et al., 2014; Langone et al., 2014; Pizza, Peterson, Baas, & Koh, 2005). This delayed expression of eMHC would indicate that post-VML there is a continued attempt at muscle regeneration which is not seen in most other acute injury muscle injury models. When there is sustained muscle regeneration, or attempted muscle regeneration, over a prolonged period of time there is the potential for MuSC depletion, as occurs in aging (Blau, Cosgrove, & Ho, 2015). This should be taken into consideration when designing therapeutics for these injuries, as it may be necessary to supplement the stem cell pool with transplanted MuSC populations.

In order for regenerating muscle to become functional, it is necessary that myofibers are innervated by the motor neuron. We were also able to characterize these regenerating fibers in terms of their neuromuscular development. Our data indicates that there is no evidence of reinnervation of the myofibers in the defect region at 14 or 28 day post-VML in an injury which is at the threshold for critical size. The fibers that are present in this area are likely a mix of myofibers that were present pre-injury as well as those which are regenerating. This is indicated in the images of injured tissue at both 14 and 28 days which shows fragmenting NMJs (class 2, Fig. 5D,F) as well as newly formed AChR clusters (class 3, Fig. S2E). Fragmenting NMJs are those which were functional pre-injury, but which are then likely denervated by the transection of the supplying motor neuron during the VML injury itself. Denervated NMJs can retain their typical morphology and the regenerating motor neuron will reinnervate at the same location with the guidance of Schwann cells between 4 and 9 days post-injury, but will then begin to display the fragmented morphology we have shown if they are not innervated in this time frame (Kang, Tian, Mikesh, Lichtman, & Thompson, 2014). Multiple new AChR clusters, however, will form on newly regenerating myofibers as they mature, secreting signaling factors to the motor neuron to direct innervation (Slater, 1982). In each of these cases, we were unable to see any evidence of the motor neuron growing towards these junctions to reinnervate myofibers, new or old, 14 or 28 days after VML injury. This could potentially be due, in part, to the destruction of guiding Schwann cells for motor neuron regeneration in VML, similarly to the loss of the guiding basal lamina for myofibers. These findings are further supported by the persistent presence of centrally located nuclei in critically sized injuries at these time points as well (Fig. 4F), as it has been shown that myonuclei will remain centrally located until the myofiber becomes functionally mature (Roman et al., 2017). This substantial dysregulation of re-innervation in the muscle regeneration process has been studied previously, both clinically (Han et al., 2016) and pre-clinically (Quarta et al., 2017), indicating similar results and which also implicate the potential for post-injury physical rehabilitation to initiate re-innervation both before and after surgical placement of a therapeutic.

A compounding factor that needs to be taken into consideration when there is nerve trauma, subsequent denervation, and re-innervation, is the potential change in skeletal muscle fiber type. During aging, there is the loss of motor units over time which is a major contributor to muscle atrophy. Typically, during this process fast motor units will leave fast fibers denervated, some of which will be re-innervated by the sprouting of slow motor units (Faulkner, Larkin, Claflin, & Brooks, 2007). As each muscle has a unique composition and distribution of fiber types (Burkholder, Fingado, Baron, & Lieber, 1994), denervation after VML may additionally have an impact on fiber type. With this in mind, therapeutics aimed at the re-innervation process may have an impact on functionality of muscle based on a re-distribution of fiber type, which may compound functional differences between injured and control muscles.

One of the most heavily studied thrusts in tissue engineering is the vascularization of tissue engineered constructs and strategies for encouraging angiogenesis. This is clearly a necessary criterion for creating a tissue engineered construct for skeletal muscle, as it is a highly vascularized tissue. There have been several studies which have shown tissue engineered strategies which promote successful formation of vessels after VML injury (M.-T. Li et al., 2017; VanDusen, Syverud, Williams, Lee, & Larkin, 2014). Interestingly, in our critically sized VML defect with no treatment we saw increased vascular volume as compared to the contralateral control. We had originally hypothesized that if we saw sufficient re-vascularization the defect, there would have been vascular remodeling in the injured tissue to result in a vascular network that closely resembled the control by 28 days. However, this increase in vascularization may have potentially been driven by the large fibrotic response, as it is known that angiogenesis and fibroplasia go hand-in-hand in the wound healing process (Greaves, Ashcroft, Baguneid, & Bayat, 2013). These results are impactful as this indicates a critically sized VML injury does not require a decrease in the vasculature volume of the injured tissue, but in fact an increase of vessels can be present up to 28 days post-injury. Our results would indicate that encouraging vascularization does not need to be a critical focus for post-VML therapeutics as the vascularization of the injury space is natively upregulated.

Based on our results assessing the muscle stem cell niche, it would indicate that a down-regulation in the fibrotic progenitor cells and an up-regulation of neural regeneration would be most beneficial for the recovery of functional skeletal muscle after VML. It should be noted that there were some limitations in the ability to always successfully image the defect area, as can be seen in one Gomori’s Trichrome image (Fig. 3F). In this image there appears to be an area unoccupied by any tissue, however there are macrophages seen in the corresponding fluorescent image. These damaged areas, filled with fibrotic tissue and likely some fatty infiltrate, were more easily damaged during cryosectioning than the surrounding muscle. When evaluating therapeutics, careful treatment of samples will be required to see subtle differences between groups. Additionally, it is important to note that while measuring muscle mass can be a useful measure when assessing muscle recovery post-VML, our results have shown there is also an increased level of fibrotic tissue after VML which would subsequently increase the measured mass of dissected muscles. For example, while the mass of the 3 mm injured samples was insignificantly different from the contralateral control, it is possible that this increase in mass can be attributed, in part, to the deposition of fibrotic tissue and therefore recovery of muscle mass is not always an indicator of improved regeneration. In future studies, it will also be necessary to consider the temporal component of regeneration. As our results indicate a delayed and subsequently sustained myofiber regeneration pattern, it is possible that varying the treatment time points will have an impact on the ability to create a pro-regenerative environment which encourages functional healing.

## Conclusion

A 3 mm full thickness defect to the mouse quadriceps, constituting approximately 15% of the muscle mass, can be used as a model for the transition point from a sub-critical to critically sized VML defect. This transition point can be characterized as a point of non-bridging of damaged myofibers into the injury space, as seen by persistent fibrotic tissue as well as chronic inflammatory response. While significant myofiber regeneration was seen, indicated by eMHC^+^ myofibers as well as myofibers with centrally located nuclei, there was no evidence of re-innervation at 14 or 28 days post-injury. The lack of re-innervation by the motor neuron into the injury space is an additional characteristic of the critical transition in VML injury sizes. However, at this stage vasculature did successfully penetrate the defect area. Overall, the regenerative responses of these muscle stem cell niche components can be used in the analysis of other VML injury models to determine the critical size for a different muscle or animal model. Additionally, this study yields insight into which areas of focus may be necessary to include in a VML therapeutic.

## Authorship

SEA, NJW, and YCJ designed the study, analyzed the data and wrote the manuscript. SEA, WMH, MM, VS, and AM conducted experiments, analyzed data, and reviewed the manuscript. MAR, CLSE, MO, and EAB provided significant contributions to methodology and data analysis and reviewed the manuscript. ES and YCJ generated and maintained animals used in this study.

## Acknowledgements

Research reported in this publication was supported by the National Institute of Arthritis and Musculoskeletal and Skin Diseases of the National Institutes of Health under Award Number R21AR072287 (YCJ). This work was conducted while Shannon E. Anderson was a Trainee on the NIH/NIGMS-sponsored Cell and Tissue Engineering (CTEng) Biotechnology Training Program (T32GM008433). The content is solely the responsibility of the authors and does not necessarily represent the official views of the National Institutes of Health. We thank the Physiological Research Laboratory and core facilities at the Parker H. Petit Institute of Bioengineering and Bioscience at the Georgia Institute of Technology for the use of shared equipment, services, and expertise.

## Figure Captions

**Supplementary Figure 1. Regenerating myofibers in 2, 3, and 4 mm injuries.** (A) Quantification of eMHC+ fibers for each injury size at 14 days post-VML. Each of the same replicate slides used for cross-sectional area quantification were used to count eMHC+ fibers. Stitched images of the entire samples were counted for eMHC+ stained fibers. Each replicate slide was counted twice. All data points shown with bars indicating mean +/− SEM (B,C) Relative frequency histograms were generated from the same data as shown in Figure 4H,I.

**Supplementary Figure 2. Neuromuscular junction imaging 14 and 28 days post-VML.** (A-D) The same images as seen in Figure 5C-F are shown here with all channels (Thy-1 is green, BTX is red, and DAPI is blue). Images from 14 days control (A) and VML (B) as well as 28 days control (C) and VML (D). A representative image is shown of 2 newly regenerating AChR clusters (indicated by orange arrowheads) in an injured sample from 14 days post-injury. Only the BTX channel is shown in red. All scale bars are 50 µm.

**Supplementary Figure 3. Pairwise comparison of vascular diameter histogram bins.** Values of vascular diameter from VML an contralateral control for each diameter bin was analyzed for significant differences between the injured and control samples. A pairwise t-test was used for each. The bins shown (A-H) had *p*<0.1, where *p*<0.05 was considered significant. * *p*<0.05, ** *p*<0.01, \*\*\*\**p*<0.0001.

**Supplementary Table 1.** Descriptive statistics of binned values in cross-sectional area histograms from 14 and 28 day timepoints.

| **14 days post-VML** |  |  |  |  |
| --- | --- | --- | --- | --- |
|  | **Control** | **2 mm** | **3 mm** | **4 mm** |
| **Total Number of Values** | 31369 | 26650 | 21782 | 13478 |
| **Median ($\mu\text{m}^2$)** | 2139.28 | 1601.23 | 1446.04 | 959.14 |
| **Mean ($\mu\text{m}^2$)** | 2174.41 | 1908.64 | 1805.89 | 1490.62 |
| **Std. Deviation ($\mu\text{m}^2$)** | 1283.41 | 1441.70 | 1472.19 | 1391.08 |
| **Std. Error of Mean ($\mu\text{m}^2$)** | 7.25 | 8.83 | 9.98 | 11.98 |
| **28 days post-VML** |  |  |  |  |
| **Total Number of Values** | 32926 | 49567 | 43347 | 29022 |
| **Median ($\mu\text{m}^2$)** | 1801.70 | 1504.63 | 1304.01 | 939.43 |
| **Mean ($\mu\text{m}^2$)** | 1982.01 | 1705.67 | 1586.81 | 1357.27 |
| **Std. Deviation ($\mu\text{m}^2$)** | 1231.77 | 1165.05 | 1194.69 | 1237.08 |
| **Std. Error of Mean ($\mu\text{m}^2$)** | 6.79 | 5.23 | 5.74 | 7.26 |

